# Distinct spatiotemporal immunometabolic remodeling following acute traumatic brain injury

**DOI:** 10.64898/2026.07.28.740519

**Authors:** Zohreh Erfani, Erin H. Seeley, Erik J. Plautz, Brenda L. Bartnik-Olson, Jae Mo Park

## Abstract

Traumatic brain injury induces profound metabolic reprogramming across neurons, astrocytes, and microglia, yet the spatiotemporal organization of these metabolic responses remains poorly understood. Because pyruvate is uniquely positioned in cerebral metabolism by connecting glycolysis, lactate metabolism, and the tricarboxylic acid cycle, we combined matrix-assisted laser desorption/ionization (MALDI) mass spectrometry imaging with *in vivo* administration of isotopically labeled pyruvate and immunohistochemistry to characterize cell-type-wise metabolic remodeling in a controlled cortical impact rat model during the acute and subacute phases of injury. TBI induced distinct spatiotemporal immunometabolic remodeling across neuroglial compartments. Microglia- and macrophage-enriched regions exhibited selective accumulation of citrate, succinate, and itaconate, consistent with inflammatory metabolic rewiring. In contrast, astrocyte enriched regions showed increased glutamine and malate abundance, indicative of altered neuron-astrocyte metabolic coupling such as remodeling of the glutamate-glutamine cycle and enhanced anaplerotic metabolism. These metabolic signatures evolved with distinct regional and temporal distributions, identifying compartmentalized metabolic responses. Notably, labeled isotopologues of selected metabolites, including glutamate and citrate derived from administered pyruvate, changed before the corresponding metabolite pools. Together, these findings describe the spatiotemporal landscape of immunometabolic remodeling following acute TBI, uncover metabolically distinct microglial/macrophage and astrocytic responses during secondary brain injury, and identify candidate metabolic pathways for therapeutic intervention and metabolic imaging.

## INTRODUCTION

Traumatic brain injury (TBI) is a leading cause of death and life-long disability, with an estimated annual incidence of 64 to 74 million cases worldwide [29, 40]. TBI progresses through two distinct phases: an initial mechanical insult followed by a complex cascade of secondary metabolic injury. The primary injury results directly from impact, leading to intracranial hemorrhage, contusions, cellular damage, and vascular disruption [27]. This sets the stage for secondary injury, which is characterized by metabolic dysfunction and neuroinflammation [13]. The acute to subacute phases of secondary injury is particularly informative, as it encompasses intense neuroinflammatory activity, cellular stress, and bioenergetic remodeling, yet precedes irreversible tissue damage [12]. Therefore, this phase represents a critical therapeutic window during which metabolic interventions may mitigate secondary brain injury.

Neuroinflammatory responses are orchestrated by a diverse array of cells and signaling molecules. In the early stages, brain-resident microglia are activated, and peripheral neutrophils are recruited, succeeded by infiltration of lymphocytes and monocyte-derived macrophages [1, 7]. This cellular response is modulated by a dynamic interplay of pro- and anti-inflammatory cytokines that govern both the initiation and resolution of inflammation [26]. Astrocytes, essential regulators of CNS homeostasis, undergo reactive astrogliosis, a process involving both morphological and functional changes [34]. Through the release of cytokines and chemokines, these glial cells further contribute to immune cell recruitment and activation [6, 9, 46, 55]. The consequences of neuroinflammation can persist indefinitely, influenced by biological factors in demographics, injury types, and therapeutic interventions [31].

Neurons, astrocytes, and microglia exhibit distinct metabolic responses to the neuroinflammatory environment post-TBI, adjusting their bioenergetic states to meet altered functional demands. Characterizing these metabolic alterations is essential for understanding the cellular mechanisms underlying secondary injury and may guide therapeutic strategies and long-term care. However, cell-type-specific immunometabolic changes during secondary injury remain poorly defined. To address this gap, *in situ* and ideally *in vivo* assessments of metabolism are required.

Pyruvate is uniquely well-suited as a tracer for evaluating central carbon metabolism. Positioned at the junction of glycolysis, lactate metabolism, and the tricarboxylic acid (TCA) cycle, pyruvate integrates signals from glucose and lactate pathways. Pyruvate’s conversion to lactate, alanine, oxaloacetate, or acetyl-CoA enables simultaneous assessment of its metabolic fate towards cytosolic and mitochondrial fluxes. Compared to upstream tracers like glucose, pyruvate bypasses multiple regulatory enzymatic steps, providing more direct insights into mitochondrial metabolism and yielding clearer isotopomer distributions.

In this study, we investigate the spatiotemporal evolution of cerebral metabolism from acute to subacute phases following TBI using matrix-assisted laser desorption ionization (MALDI) mass spectrometry imaging (MSI) in a rat TBI model. This approach enables spatial mapping of metabolites directly within tissue sections while preserving anatomical context. By integrating these metabolic profiles with MALDI immunohistochemistry (IHC), metabolite distributions can be interpreted alongside cell-type-specific protein markers [41]. This enables the investigation of distinct metabolic adaptations across neurons, astrocytes, and microglia that cannot be captured by whole tissue metabolomics approaches [5, 32, 36, 45]. In addition, ^13^C-labeled metabolites, produced from bolus injected [U-^13^C_3_]pyruvate, are analyzed to assess metabolic pathway activity and carbon flux [17, 23].

## MATERIALS and METHODS

### Animal Preparation and TBI Model

All animal procedures followed the Guide for Care and Use of Laboratory Animals of US National Research Council and were approved by the University of Texas Southwestern Medical Center Institutional Animal Care and Use Committee (Protocol #:2017-101802). All animal experiments are in accordance with the Animal Research: Reporting In Vivo Experiments (ARRIVE) guidelines.

A controlled cortical impact (CCI) rat model was used in this study [20]. With precise biomechanical control over impact parameters, the CCI model facilitates the correlation of measurable loading parameters with morphological and functional outcomes [49]. Twenty healthy male Wistar rats (200-250 g) were anesthetized with isoflurane (induction: 3.5-4.5%, maintenance: 2-2.5%, in a 70% nitrous oxide, 30 % oxygen carrier mix). Each rat was mounted in a stereotaxic device (David Kopf Instruments) with the head secured via nose mask and ear bars. A midline scalp incision was made to expose skull suture landmarks (i.e., Bregma and Lambda), and a 0.4 cm (ML) × 0.5 cm (AP) craniectomy was performed over the right parietal cortex, leaving the dura intact. Contralateral uninjured cortex (left) was compared with the injured region (right). A commercial CCI apparatus (Leica MyNeuroLab Impact One™ stereotaxic impactor, Leica Biosystems Richmond Inc.) was used to produce the brain injury with the following parameters: 3.0 mm diameter impactor tip-oriented perpendicular to the cortical surface, 4.4 m/s impact velocity, 1.0 mm impact depth, and 100 ms impact duration. This procedure reliably induced mild-to-moderate TBI. The craniectomy was repaired by re-securing the bone flap with dental acrylic, leaving a small hole laterally for potential fluid drainage, and the scalp incision closed with suture.

During the surgical procedure a body temperature of 37 °C was maintained using a homeothermic heating system (RightTemp, Kent Scientific Corp.). Respiration rate was visually monitored. Lidocaine (2% solution, topical at incision), Burprenorphine ER (0.6 mg/kg, s.q.), carprofen (5 mg/kg, s.q.), and saline (0.9% solution, 0.5 ml, s.q.) were given for analgesia and fluid replacement. Rats recovered in a temperature-controlled chamber (30 °C ambient air). Postoperatively, animals were regularly monitored for symptoms such as seizures, loss of consciousness, nasal bleeding, and hemorrhage. The manifestation of any of these symptoms would have prompted the early exclusion of rats from the study through euthanasia. However, none of the rats exhibited any such indications. Four surgeries were performed each day with one rat from each group to minimize potential confounders.

### Injection of [U-^13^C_3_]Pyruvate and Tissue Collection

The animals were under observation following surgery and were randomly assigned to different post-surgery timepoint subgroups. The first group was sacrificed 2 hours post-surgery (n = 5), second group 2 days post-surgery (n = 5), third group 5 days post-surgery (n = 5) and the last group 10 days post-surgery (n = 5). One brain sample from the 2-hour post-surgery group was not analyzed because the tissue was damaged (shredded) during shipment. Each rat was administered with a bolus of [U-^13^C_3_]pyruvate (0.75 mmol/kg body weight, injection rate = 0.25 mL/s). 67.5 mg of sodium [U-^13^C_3_]pyruvate was dissolved to 5-mL water to make 120-mM pyruvate. The brain was rapidly dissected within 2 minutes following decapitation and immediately immersed in liquid nitrogen to be fixed. The bolus injection was to capture early metabolic incorporation while minimizing extensive isotopic redistribution across cellular compartments. Short labeling windows are commonly used in ^13^C metabolic studies to probe the initial routing of substrate carbon rather than steady-state labeling patterns since longer labeling periods can increase isotopic equilibration and obscure early metabolic pathways of tracer incorporation. Previous studies using bolus administration of ^13^C-labeled substrates have shown that even short tracer exposure can reveal distinct neuronal and astrocytic metabolic fluxes and changes in key metabolites [18, 19, 47].

### MALDI MSI

Brains were sectioned at 12 μm thickness within in the TBI area and collected onto mass spectrometry-compatible slides. Serial sections were collected for H&E staining. Sections were coated with 7 mg/mL N-(1-naphthyl) ethylenediamine dihydrochloride in 70% methanol using an HTX M5 Robotic Reagent Sprayer over 8 passes with the following parameters; a flow rate of 0.120 mL/min, a track speed of 1200 mm/min, a track spacing of 2 mm, a crisscross track pattern, and a nozzle temperature of 75 °C. Sections were imaged on a timsTOF fleX MALDI QTOF mass spectrometer (Bruker, Billerica, MA, USA) in negative ion mode at 100 μm spatial resolution over the *m/z* range 50-600 with 1000 laser shots summed per pixel. Instrument parameters were as follows; a Funnel 1 radiofrequency (RF) of 75.0 peak-to-peak voltage (V_pp_), a Funnel 2 RF of 100.0 V_pp_, a Multipole RF of 150.0 V_pp_, a Collision Energy of 5.0 eV, a Collsion RF of 500.0 V_pp_, a Quadrupole Ion Energy of 5.0 eV, a Transfer Time of 40 μs, and a Pre Pulse Storage of 3.0 μs.

A peak list of target metabolite along with their predicted ^13^C isotopes was imported into SCiLS Lab (Bruker), and images were generated for each *m/z*. Predicted abundances of each isotope were generated using the Bruker Isotope Pattern. The intensity of each isotope from the targeted metabolite list was exported to Excel (Microsoft). The abundance of each isotope as a percentage of the monoisotopic peak was calculated and compared the predicted natural abundance.

### MALDI IHC

MALDI IHC was also acquired from two rats in each group (n = 8) according to standard AmberGen’s (Watertown, MA, USA) standard protocol [50]. Briefly, the sections were fixed with paraformaldahyde before rehydration and antigen retrieval with Tris-EDTA at 95 °C for 30 minutes. Sections were treated with blocking buffer for 1 hour before incubation with NeuN, glial fibrillary acidic protein (GFAP), and vimentin (VIM) for 1 hour at 37°C. After washing with tris-buffered saline and ammonium bicarbonate, the sections were photocleaved with UV light and then coated with α-cyano-4-hydroxycinnamic acid matrix. Images were acquired on the timsTOF fleX in positive ion mode at 100 μm spatial resolution over the *m/z* range 800-180 with 1000 laser shots summed per pixel. Instrument parameters were as follows; a Funnel 1 RF of 250.0 V_pp_, a Funnel 2 RF of 500.0 V_pp_, a Multipole RF of 5000.0 V_pp_, a Collision Energy of 25.0 eV, a Collsion RF of 3000.0 V_pp_, a Quadrupole Ion Energy of 25.0 eV, a Transfer Time of 110 μs, and a Pre Pulse Storage of 15.0 μs. Images of the reporter peptides were visualized using SCiLS Lab.

### Image Analysis and Statistical Analysis

Group allocation was blinded during MALDI data collection and postprocessing until statistical analysis, except for Z.E. MALDI MSI and MALDI IHC images were imported into MATLAB (R2024b; MathWorks Inc., Natick, MA, USA) for quantitative analysis. Injured brain regions were identified based on hyperintense (glutamine, malate, citrate, itaconate, succinate) or hypointense (glutamate, aspartate) signals, using intensity thresholds derived from manually selected contralateral, normal-appearing regions. The outermost contour of the thresholder area was used to define region-of-interest (ROI) boundaries for lesion size measurement and subsequent metabolite quantification. Within each ROI and its contralateral counterpart, intensities of unlabeled and labeled isotopologues were averaged. Fractional enrichment was calculated as the ratio of the isotopologue of interest to the total signal across all isotopologues. The contralateral hemisphere served as an internal reference region for comparison. This approach enables paired comparisons within the same animal and reduces inter-animal variability.

Paired t-tests (α = 0.05, two-tailed) were used to assess differences between injured and contralateral regions. For spatial correlation analysis, MALDI IHC images were co-registered to MSI data via rigid translation and rotation using MATLAB. Metabolite ratio maps were generated by pixel-by-pixel division of corresponding metabolite images. All data are reported mean ± standard deviation.

To characterize temporal trends in metabolite intensities and lesion sizes, mean values from hyper- or hypointense ROIs were normalized to contralateral values and fitted to a spline curve using Prism 10 (version 10.5.0; GraphPad Software, San Diego, CA, USA). All figures were prepared using Prism 10 (charts), MATLAB (linear regressions), and Affinity Designer (v1.10.8; Serif, West Bridgford, UK).

## RESULTS

For metabolic profiling of acute and subacute conditions, CCI-induced TBI rats were analyzed at 2 h, 2, 5, and 10 days post-injury. Two minutes prior to tissue collection, each rat received a bolus of [U-^13^C_3_]pyruvate. Spatial metabolomics analysis was performed using MALDI MSI, as outlined in **Figure 1**. Both unlabeled and labeled metabolites were detected in the brain, highlighting mitochondrial metabolism through pyruvate dehydrogenase (PDH) and pyruvate carboxylase (PC). Lactate signals were generally weak and variable across samples due to poor ionization efficiency of lactate in negative ion mode. Overall, PDH-derived M+2 isotopologues were more abundant than PC-derived M+3 isotopologues after bolus injection of [U-^13^C_3_]pyruvate, consistent with a previous NMR and mass spectrometry study [8].

**Figure 1.**
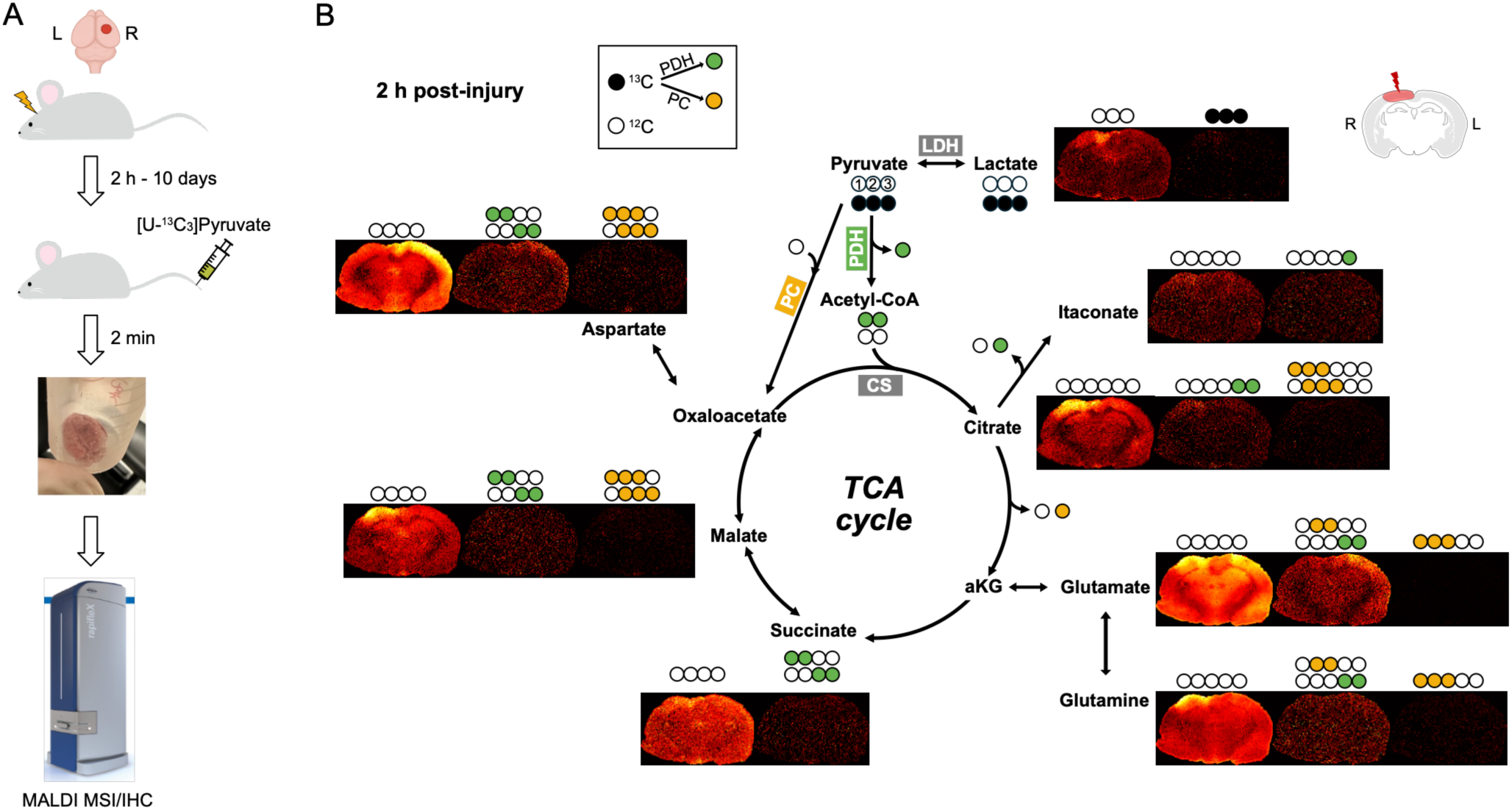
Study overview. (A) Schematic of the experimental procedure. Brain tissues were collected from four groups of CCI rats at 2 h, 2, 5, and 10 days post-injury. The CCI was performed in the right hemisphere. Tissue harvesting occurred 2 min after a bolus injection of [U-^13^C_3_]pyruvate for MALDI MSI and IHC. (B) Diagram illustrating the labeled carbons in [U-^13^C_3_]pyruvate into the TCA cycle via PDH and PC. Filled and open circles represent labeled (^13^C) and unlabeled (^12^C) carbons, respectively. Color-coded circles indicate ^13^C incorporation via PDH (green) or PC (yellow). Representative MSI images from a rat brain at 2 h post-injury display both intrinsic (unlabeled) and exogenously derived (^13^C-labeled) metabolites, including lactate, citrate, itaconate, glutamate, glutamine, succinate, malate, and aspartate.

### Enhanced Citrate Reflects Microglial and Macrophage Activation

Citrate level was significantly elevated in the injured region as showcased in two CCI rats at 5 days post-injury (**Figure 2A**). The distribution of citrate signals was positively correlated with VIM (ρ = 0.276, *P* < 0.0001) and negative correlation with GFAP (ρ = −0.384, *P* < 0.0001) and NeuN (ρ = −0.257, *P* < 0.0001), suggesting a microglial and macrophage origin (**Figure 2B**) [4, 15, 25, 51]. Citrate levels were consistently elevated in the injured cortex as compared with the contralateral normal-appearing brain region across all timepoints (*P* = 0.042 at 2 h, 0.043 at 2 days, 0.039 at 5 days, and 0.001 at 10 days), with the greatest increase at 5 days post-injury (**Figure 2C-D**). In contrast, the size of the citrate-enhanced region peaked immediately after injury, followed by monotonic decrease, implying that global-to-local and gradually more intense microglial and macrophage activation during day 0 – 5 (**Figure 2E**).

**Figure 2.**
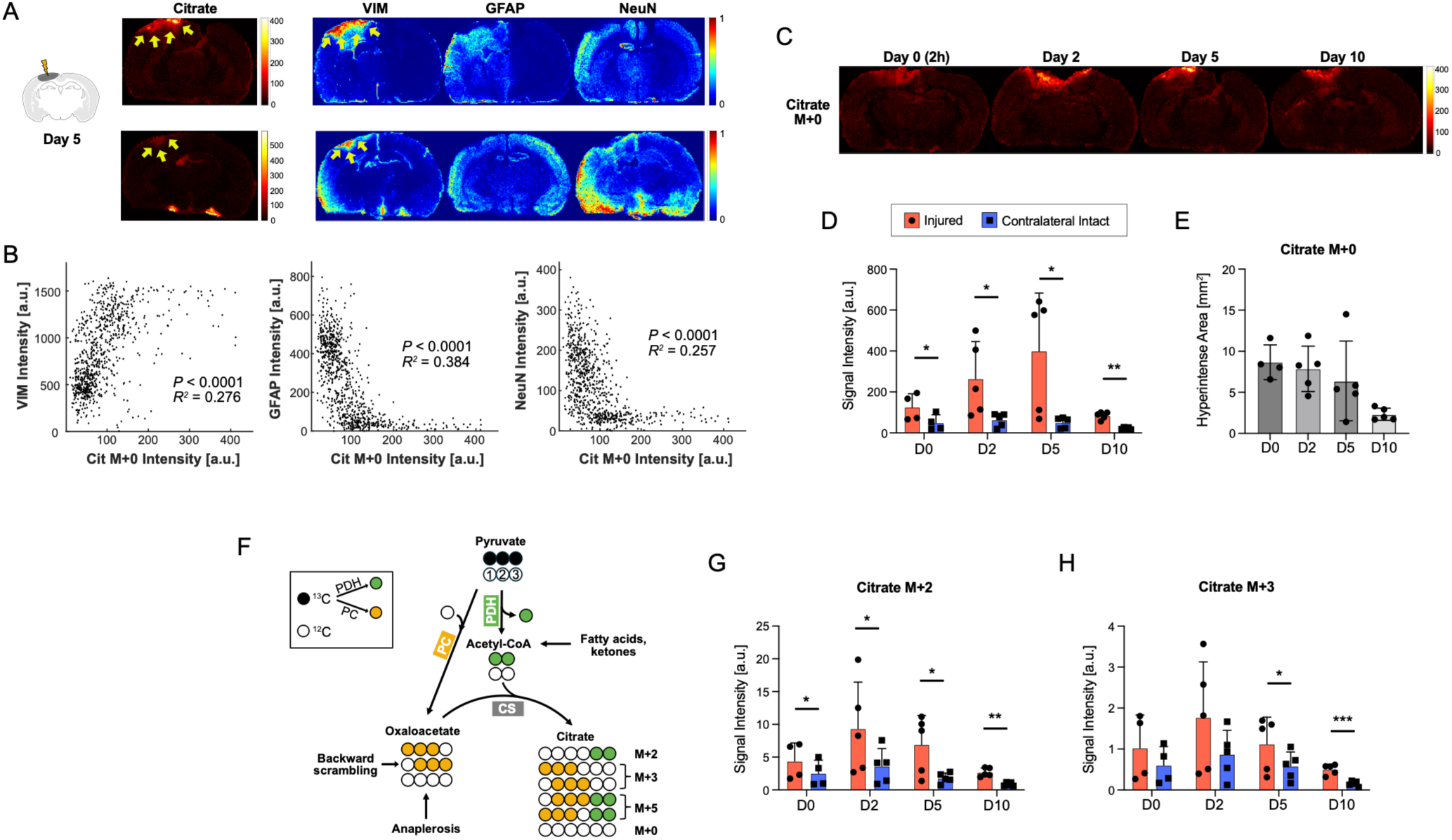
Enhanced citrate levels in microglia- and macrophage-enriched regions following TBI. (A) MALDI MSI of unlabeled citrate (M+0) from two representative rats at 5 days post-injury, together with matched MALDI IHC for VIM, GFAP, and NeuN. (B) Scatter plots comparing M+0 citrate intensity with VIM, GFAP, and NeuN signals, demonstrating a positive correlation with VIM and negative correlations with GFAP and NeuN. (C) Spatial distributions of unlabeled citrate (M+0) from representative rats at 2 h (day 0), 2, 5, and 10 days post-injury. (D) Quantification of M+0 citrate intensity in the injured and contralateral (intact) hemispheres, showing sustained citrate elevation in the injured cortex, with the greatest difference observed at day 5. (E) Quantification of the citrate-hyperintense area over time. (F) Schematic illustrating the generation of citrate isotopologues via PDH (M+2) and PC (M+3) pathways, as well as possible contributions from anaplerotic pathways, fatty acid oxidation, and ketone metabolism. (G, H) Signal intensities of M+2 and M+3 citrate over time, demonstrating increased labeling in the injured brain relative to the contralateral side.

In addition to unlabeled citrate, two labeled citrate isotopologues were detected due to [U-^13^C_3_]pyruvate bolus: M+2 citrate, generated from [1,2-^13^C_2_]acetyl-CoA via PDH, and M+3 citrate, generated from [1,2,3-^13^C_3_]oxaloacetate via PC (**Figure 2F**). M+2 citrate was elevated at all timepoints (*P* = 0.038, 0.049, 0.039, 0.004), peaking at 2 days after injury (**Figure 2G**). Notably, the earlier increase in M+2 citrate relative to unlabeled citrate suggests that changes in metabolic flux precede expansion of the citrate pool. M+3 citrate was significantly elevated only at 5 and 10 days (*P* = 0.035 and 0.001; **Figure 2H**). Despite these increases, the fractional enrichment of M+2 and M+3 citrate was reduced in the injured cortex, suggesting dilution by an expanding unlabeled citrate pool (**Figure S1**).

### Itaconate and Succinate Defines Immunometabolic Phenotypes Following TBI

Consistent with inflammatory immunometabolic remodeling, unlabeled (M+0) itaconate and succinate accumulated in the injured cortex following TBI (**Figure 3**). Both metabolites increased as early as 2 h after injury, peaked at day 2 and day 5, and declined by day 10 (itaconate: *P* = 0.023, 0.019, 0.029, and 0.014; succinate: *P* = 0.019, 0.042, 0.094, and 0.016). Spatial correlation analysis demonstrated that both metabolites negatively correlated with NeuN (itaconate: ρ = −0.173, *P* < 0.0001; succinate: ρ = −0.151, *P* < 0.0001) and GFAP (itaconate: ρ = −0.223, *P* < 0.0001; succinate: ρ = −0.198, *P* < 0.0001), while showing positive spatial association with VIM (itaconate: ρ = 0.166, *P* < 0.0001; succinate: ρ = 0.055, *P* = 0.10), consistent with enrichment in microglia-/macrophage-associated regions (**Figure S2**).

**Figure 3.**
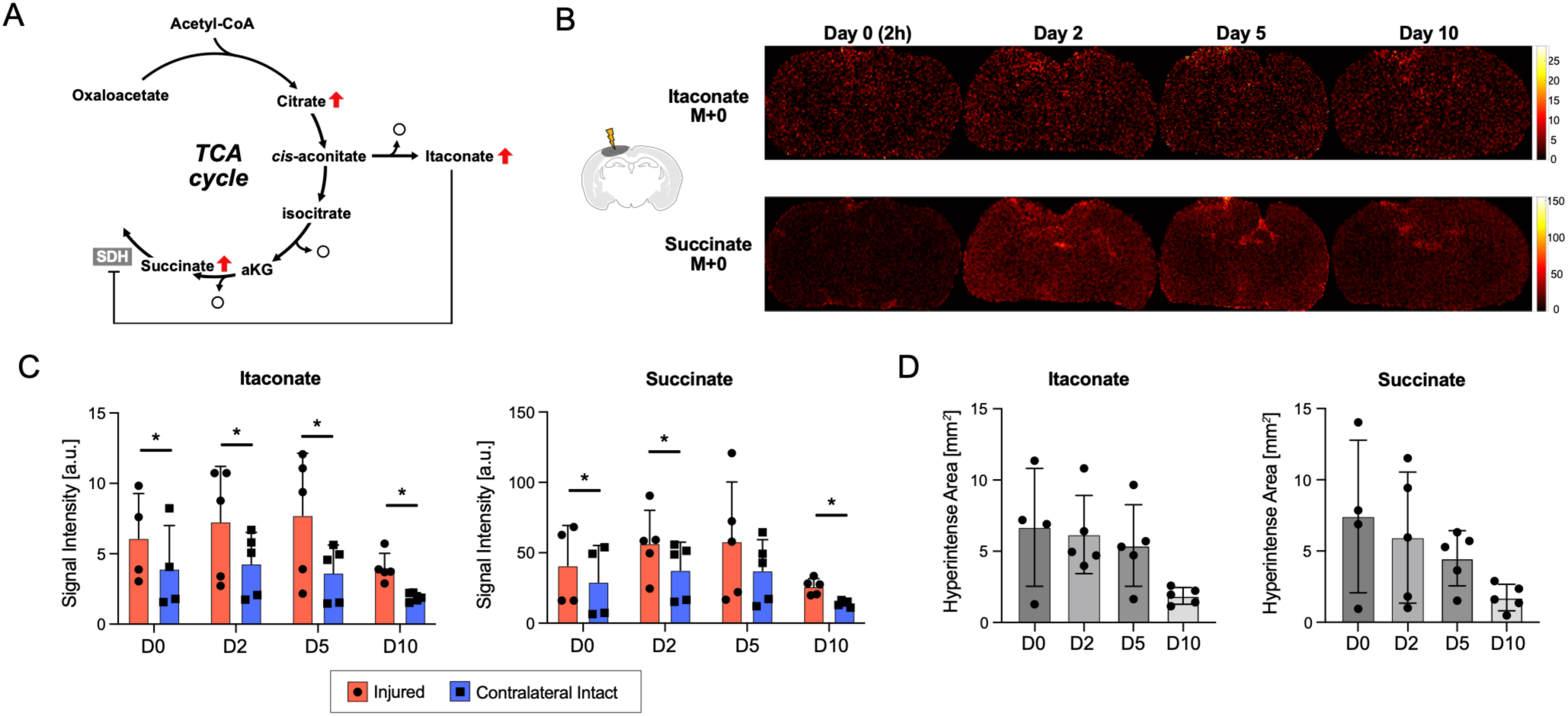
Itaconate and succinate accumulation in injured brain regions. (A) Metabolic responses to inflammation elevate the synthetic pathway to itaconate (via *cis*-aconitate), subsequently suppressing succinate dehydrogenase (SDH) and resulting in succinate accumulation. (B) MALDI MSI showing spatial distributions of unlabeled (M+0) itaconate and succinate at 2 h (day 0), 2, 5, and 10 days post-injury. (C) Quantification of M+0 itaconate and M+0 succinate showed elevated levels in the injured cortex at 2 h - 5 days post-injury, followed by a decline at day 10. (D) Quantification of the hyperintense region size based on itaconate and succinate M+0 maps across time points.

In contrast to the sustained elevation of the unlabeled metabolite pools, stable-isotope tracing showed earlier and more transient changes in metabolic flux (**Figure S3**). M+1 itaconate ([1-^13^C]) and M+2 succinate ([1,2-^13^C_2_] and [3,4-^13^C_2_]) both peaked at day 2 and declined rapidly thereafter, preceding the prolonged accumulation of their corresponding unlabeled metabolites. Although the increase in M+1 itaconate did not reach statistical significance (*P* = 0.14 at day 5, *P* = 0.11 at day 10), the advanced temporal profile of both labeled isotopologues suggests that alterations in immunometabolic flux occur before expansion of the endogenous metabolite pools.

### Distinct Spatiotemporal Remodeling of Glutamate and Glutamine Metabolism Following TBI

M+0 glutamate and glutamine showed opposing spatiotemporal responses following TBI, consistent with disrupted metabolic coupling between neurons and astrocytes (**Figure 4**). Unlabeled glutamate remained largely unchanged at 2 h after injury but was significantly reduced from day 2 onwards (*P* = 0.34, 0.026, 0.038, and 0.0002), reflecting progressive neuronal dysfunction or loss. In contrast, M+0 glutamine increased as early as 2 h after injury and remained elevated throughout the study period (*P* = 0.041, 0.034, 0.017, and 0.0008). Spatially, glutamine accumulated predominantly in tissue surrounding glutamate-depleted regions, indicating astrocytic response to injury. The glutamate-hypointense area and glutamine-hyperintense area expanded at days 2 and 5 before partially resolved by day 10.

**Figure 4.**
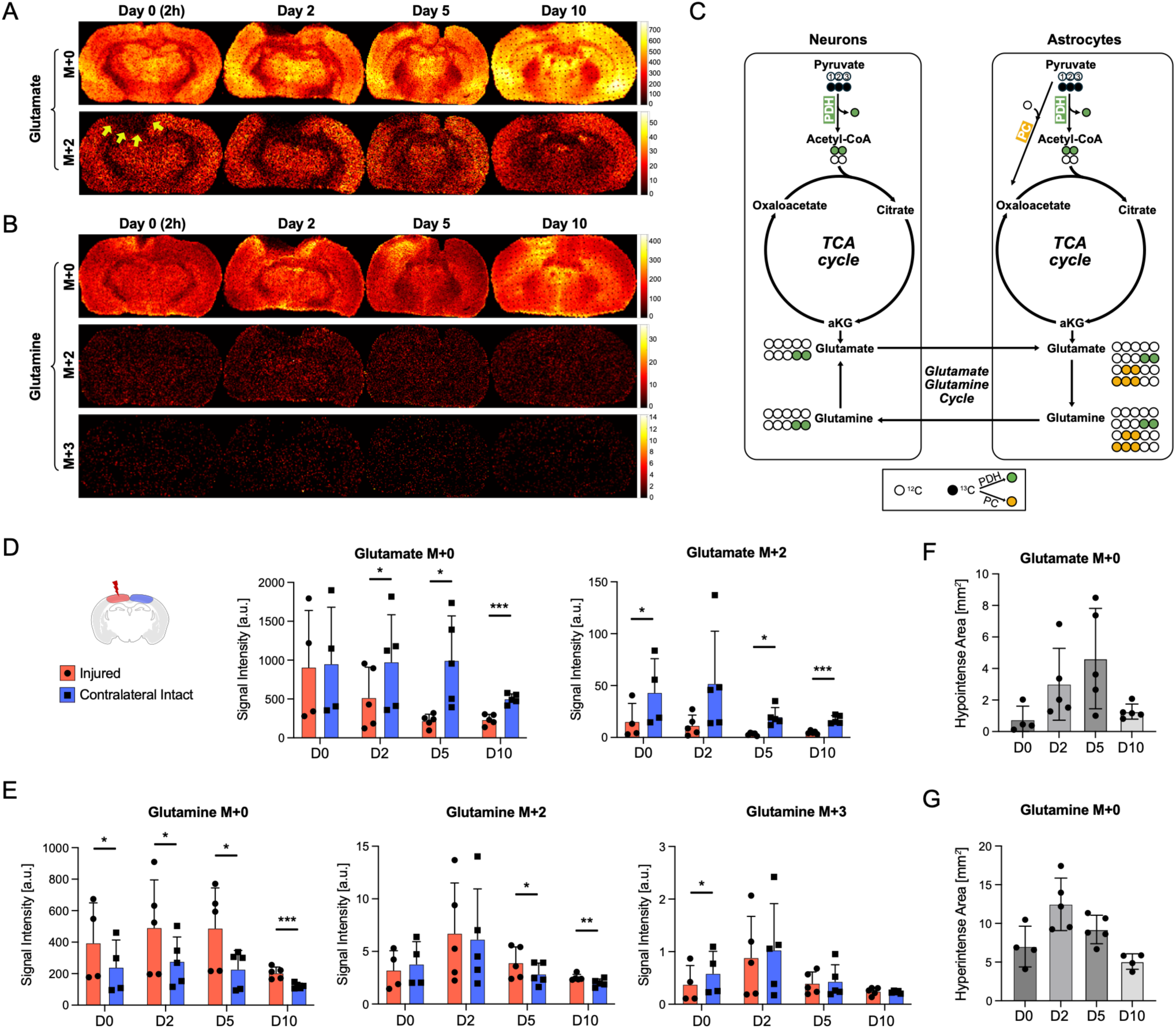
Spatiotemporal remodeling of glutamate and glutamine metabolism after TBI. (A) MALDI MSI showing spatial distributions of unlabeled (M+0) and PDH-derived (M+2) glutamate at 2 h, 2, 5, and 10 days post-injury. (B) MALDI MSI of glutamine depicting spatial distributions of unlabeled (M+0), PDH-derived (M+2), and PC-derived (M+3) isotopologues across time points. (C) Schematic summarizing the cell-type-specific origins of glutamate and glutamine isotopologues. In neurons, pyruvate is oxidized via PDH to acetyl-CoA, generating M+2 glutamate, which can be transferred to astrocytes and converted to M+2 glutamine via the glutamate-glutamine cycle. In astrocytes, PC-mediated carboxylation of pyruvate produces M+3 oxaloacetate, which combines with unlabeled acetyl-CoA to form M+3 glutamate and glutamine. (D) M+0 glutamate was not altered at 2 h post-injury, then progressively declined in the injured cortex over time. In contrast, M+2 glutamate was instantly reduced in the injured cortex and the reduction persisted at all time points, reflecting acute disruption of neuronal oxidative metabolism and diminished pyruvate entry into the TCA cycle. (E) M+0 glutamine intensity increased in the injured hemisphere, indicating expansion of the glutamine pool. M+2 and M+3 isotopologues of glutamine increased in both hemispheres at 2 days post-injury. Temporal evolution of spatial abnormalities, as reflected by the size of (F) the hypointense region for M+0 glutamate and (G) the hyperintense region for M+0 glutamine.

Stable-isotope tracing of glutamate could also detect metabolic alterations that preceded changes in metabolite abundance. M+2 glutamate ([4,5-^13^C_2_] and [2,3-^13^C_2_]) decreased at 2 h after injury (*P* = 0.047), preceding the decline in the unlabeled glutamate pool, and remained decreased through day 10 (*P* = 0.091, 0.018, and 0.0005). In contrast, the temporal profiles of M+2 and M+3 glutamine largely paralleled those of unlabeled glutamine. M+2 glutamine appeared increased at day 2 in both injured and contralateral cortices. The elevated M+2 glutamine decayed slower in the injured cortex than the contralateral region, making the difference significant at days 5 and 10 (*P* = 0.018 and 0.008). Fractional enrichments of both M+2 and M+3 glutamine were consistently lower in the injured cortex, indicating diminished incorporation of pyruvate-derived carbon into glutamine despite expansion of the glutamine pool (**Figure S4**).

### Altered Malate-Aspartate Metabolism Implies Disrupted Neuron-Glia Coupling

Malate and aspartate also showed opposing metabolic responses to TBI, further supporting disruption of neuron-glia coupling (**Figure 5**). M+0 aspartate decreased in the injured cortex at all timepoints (*P* = 0.025, 0.004, 0.046, 0.0008), while M+0 malate increased (*P* = 0.020, 0.051, 0.013, and 0.004). Spatial correlation analysis demonstrated that M+0 aspartate was strongly associated with NeuN (ρ = 0.605, *P* < 0.0001) and negatively with GFAP and VIM (**Figure S5**), consistent with a predominantly neuronal distribution. In contrast, M+0 malate showed positive correlations with both GFAP (ρ = 0.101, *P* < 0.0001) and VIM (ρ = 0.201, *P* < 0.0001), indicating preferential accumulation within glia-enriched regions.

**Figure 5.**
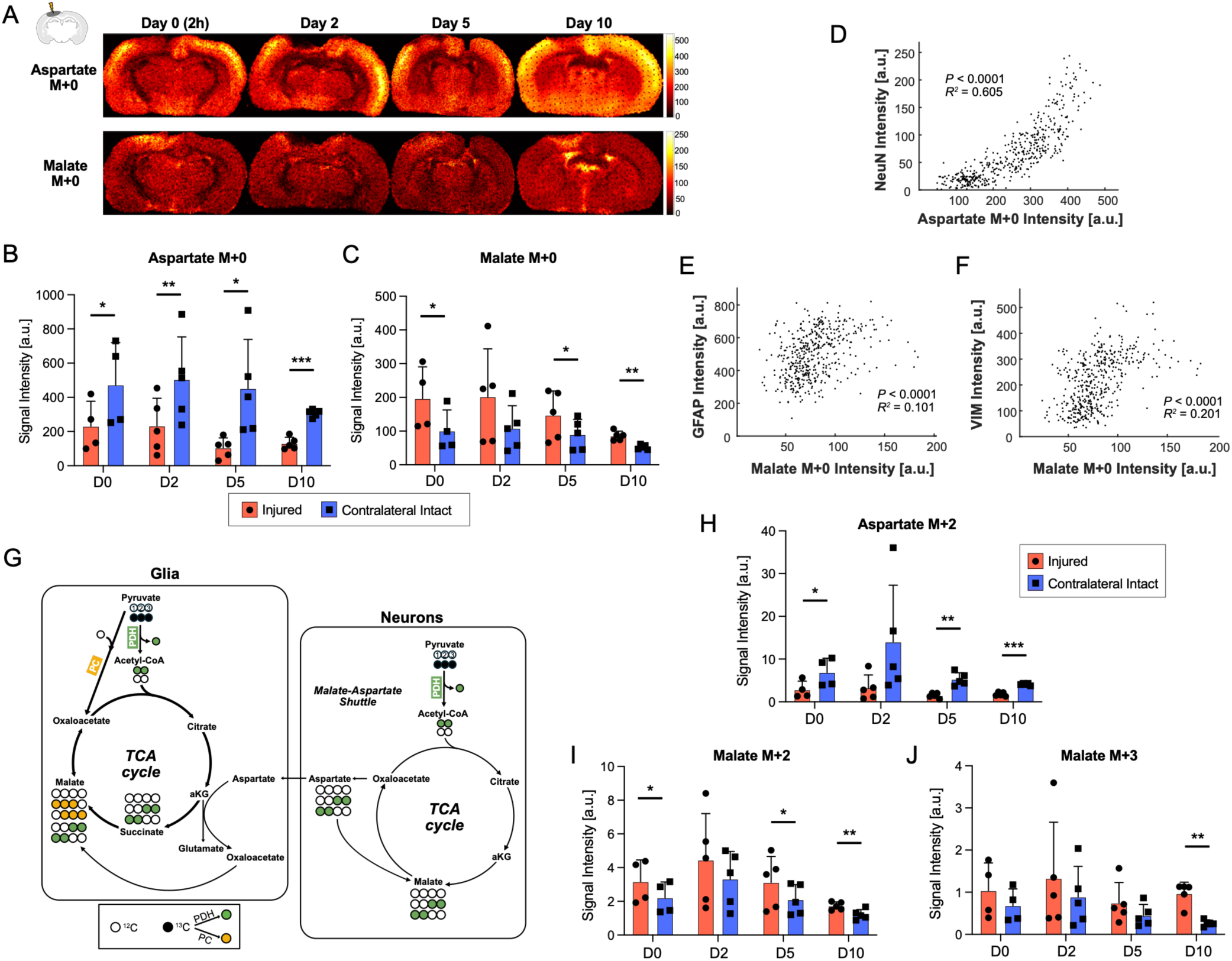
Disrupted neuron-glia malate-aspartate metabolism by TBI. (A) MALDI MSI showing the spatial distribution of intrinsic (M+0) aspartate and malate at 2 h, 2, 5, and 10 days post-injury. (B, C) Quantification of M+0 aspartate and malate showing persistent depletion of aspartate and accumulation of malate within the injured cortex. (D-F) Spatial correlation analyses showing strong association of aspartate with NeuN and positive associations of malate with GFAP and VIM, consistent with neuronal localization of aspartate and glial enrichment of malate. (E, F) Scatter plots of M+0 malate with GFAP and VIM demonstrate a positive association with glial markers. (G) Schematic illustrating cell-type-specific malate and aspartate metabolism. In neurons, [U-^13^C_3_]pyruvate is oxidized via PDH, generating M+2 malate and M+2 aspartate that participate in the malate-aspartate shuttle. In glia, pyruvate enters through both PDH and PC pathways, supporting oxidative and anaplerotic metabolism. (H, I) PDH-derived M+2 aspartate and M+2 malate in the injured cortex and contralateral region over time. (J) PC-derived M+3 malate over time.

Labeled aspartate and malate showed similar patterns. PDH-mediated M+2 aspartate ([1,2-^13^C_2_] and [3,4-^13^C_2_]) in the injured region was lower than the contralateral region. Despite low signal sensitivities, both M+2 malate ([1,2-^13^C_2_] and [3,4-^13^C_2_]) and M+3 malate ([1,2,3-^13^C_3_] and [3,4,5-^13^C_3_]) were higher in the injured region, suggesting activation of oxidative and anaplerotic metabolism in glia (**Figure S5**). These observations further support disruption of neuron-glia malate-aspartate metabolism during acute and sub-acute phases of TBI.

### Composite Metabolite Ratios Capture Neuron-Glia Dysfunction and Immunometabolic Remodeling

Composite metabolite ratios highlighted coordinated neuron-glia metabolic disruption and neuroinflammation following TBI (**Figure 6**). The M+0 glutamate/glutamine ratio, which reflects neuron-astrocyte coupling, was stable over time in the contralateral intact region (3.86 – 4.39) but declined in the injured cortex to 2.08 ± 0.49 at 2 h (*P* = 0.002), 1.00 ± 0.43 at 2 days (*P* < 0.0001), and 0.61 ± 0.42 at 5 days (*P* < 0.0001), partially recovering by day 10 (1.14 ± 0.26, *P* < 0.0001). Likewise, the aspartate/malate ratio, another index of neuron-glia interaction, was consistently lower in the injured cortex relative to the contralateral hemisphere (4.86 – 5.92), reaching 1.11 ± 0.21 at 2 h (*P* = 0.008), 1.19 ± 0.36 at 2 days (*P* = 0.003), 0.96 ± 0.74 at 5 days (*P* = 0.0008), 1.51 ± 0.23 at 10 days (*P* = 0.0005).

**Figure 6.**
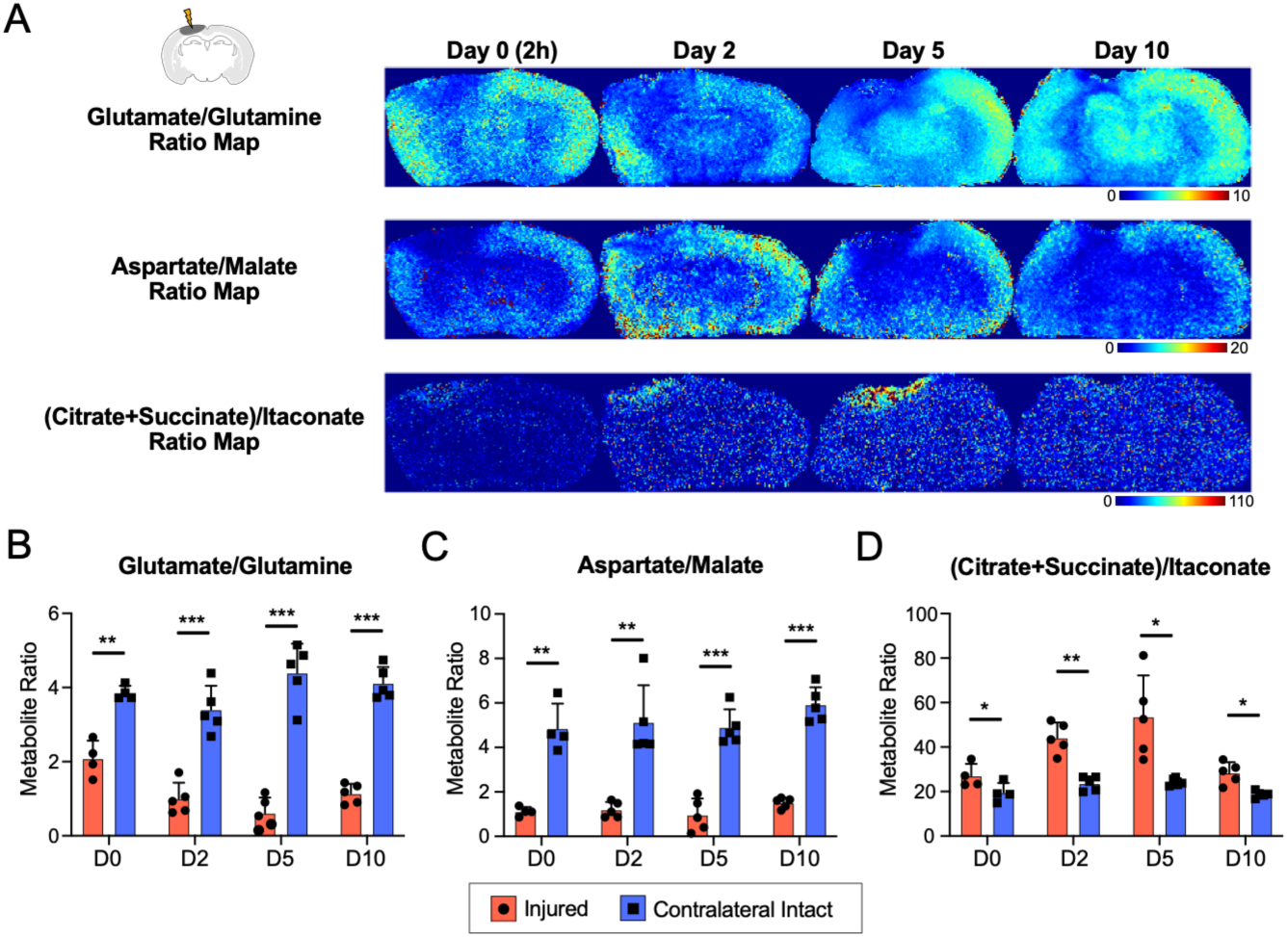
Composite metabolite ratios. (A) Based on MALDI MSI M+0, glutamate/glutamine ratio maps, aspartate/malate ratio maps, and (citrate+succinate)/itaconate ratio maps were generated at 2 h, 2, 5, and 10 days post-injury MALDI MSI. (B) Quantification of the M+0 glutamate/glutamine ratio revealed a significant decrease in the injured cortex as early as 2 h post-injury, reaching a nadir at 5 days, followed by partial recovery by day 10. (C) Quantification of the M+0 aspartate/malate ratio in the injured cortex demonstrated immediate and sustained reductions relative to the contralateral hemisphere. (D) A composite inflammatory ratio map calculated as (succinate+citrate)/itaconate, reflecting the balance between inflammatory metabolic activation and endogenous immunoregulation.

The inflammation-associated metabolite ratio, M+0 (succinate + citrate)/itaconate ratio, reflecting the relative abundance of pro- and anti-inflammatory TCA intermediates and the immunoregulatory metabolite itaconate [33], was consistently elevated in the injured cortex compared with the contralateral hemisphere (*P* = 0.031, 0.007, 0.027, and 0.013). This ratio displayed an inverse temporal pattern relative to the glutamate/glutamine and aspartate/malate ratios, consistent with progressive immunometabolic remodeling. The increasing (succinate+citrate)/itaconate suggests a progressive accumulation of inflammatory metabolites due to insufficient compensatory itaconate during 2 h – 5 days post-injury.

Across all composite metabolite ratios, the greatest contrast between the injured and contralateral cortex occurred at 5 days post-injury, suggesting that these spatially resolved metabolic indicators sensitively capture the peak of secondary injury.

## DISCUSSION

TBI alters cerebral metabolism through complex interactions among neurons, astrocytes, microglia, and infiltrating macrophages. Neurons and astrocytes coordinate neurotransmission and metabolic homeostasis, whereas microglia and macrophages serve as key immune regulators of neuroinflammatory responses during secondary injury [3]. Astrocytes regulate cerebral blood flow, buffer extracellular glutamate, and provide metabolic support to neurons [38], while activated microglia undergo extensive immunometabolic reprogramming characterized by production of metabolites such as citrate, succinate, and itaconate [42, 52]. In this study, these distinct cellular metabolic programs were reflected by metabolite-wise spatial and temporal trajectories, identifying compartmentalized metabolic remodeling across the injured brain.

Using MALDI MSI together with IHC and *in vivo* administration of [U-^13^C_3_]pyruvate, we mapped the spatiotemporal organization of metabolic remodeling during the acute-to-subacute phases following brain injury. Rather than a uniform metabolic response, individual metabolites showed distinct profiles that reflected neuroglial compartments and metabolic programs. Glutamine and malate showed relatively broad spatial expansion immediately after injury, whereas citrate, itaconate, and succinate were confined to more spatially restricted regions throughout the study period. These patterns suggest an early phase of widespread neuroglial metabolic adaptation followed by increasingly localized immunometabolic remodeling associated with activated microglial and macrophages during secondary brain injury.

Composite metabolite ratios further highlighted the disruption of neuron-glia metabolism. The marked decline in the glutamate/glutamine ratio reflected impaired metabolic interaction between neurons and astrocytes. Likewise, the reduced aspartate/malate ratio was accompanied by neuronal depletion of aspartate and glial accumulation of malate, indicating altered malate-aspartate metabolism and perturbation of metabolic neuroglial interactions. Because both the glutamate-glutamine cycle and malate-aspartate metabolism are tightly linked to mitochondrial function and neurotransmitter homeostasis, these findings support broad neuroglial metabolic reprogramming triggered by TBI. Conversely, the inflammation-associated (citrate + succinate)/itaconate ratio increased significantly after injury, peaking at 5 days in VIM-positive regions, indicating progressive immunometabolic remodeling within the inflammatory lesion.

The spatial colocalization of citrate, succinate, and itaconate accumulation is consistent with established metabolic signatures of activated microglia and macrophages. Citrate accumulation supports inflammatory activation through SLC25A1-mediated export to the cytosol, where it supports acetyl-CoA for histone acetylation, lipid biosynthesis, and further cytokine production [22, 43, 53]. Citrate-derived cis-aconitate is converted to itaconate by IRG1, providing an essential immunoregulatory mechanism through inhibition of succinate dehydrogenase (SDH), reduction of mitochondrial ROS production, and suppression of inflammatory signaling [24, 28, 48, 51, 54]. In line with these mechanisms, we observed accumulation of succinate in injured regions, suggesting that altered metabolite turnover or impaired SDH activity contributes to succinate accumulation following TBI.

Distinct alterations in glutamate, glutamine, aspartate, and malate further demonstrate disruption of neuron-astrocyte metabolic coupling. Glutamate and aspartate remained predominantly neuronal, whereas glutamine localized primarily to astrocytic regions and malate accumulated within both astrocytic- and microglial-enriched regions. Elevated glutamine together with reduced ^13^C enrichment is consistent with enhanced astrocytic uptake and detoxification of glutamate released from injured neurons. At the same time, reduced neuronal utilization of glutamine likely contributes to its regional accumulation. In addition, reciprocal changes in malate and aspartate suggest altered malate-aspartate metabolism, potentially reflecting impaired metabolic exchange between neurons and glia. Because aspartate serves as both a key component of the malate-aspartate shuttle and a precursor for glutamate synthesis, depletion of neuronal aspartate may contribute to reduced glutamate production and impaired mitochondrial metabolic homeostasis after injury. Notably, neuronal aspartate depletion occurred immediately after injury, preceding substantial loss of the glutamate pool, suggesting that disruption of aspartate-dependent metabolism may contribute to the subsequent decline in neuronal glutamate [30, 37, 44]. These observations agree with previous isotope-tracing studies reporting enhanced astrocytic glutamate conversion despite relatively preserved neuronal oxidative metabolism after TBI [2].

A unique strength of the current study is the integration of stable-isotope tracing with spatial metabolomics. Conventional metabolic studies rely on steady-state isotope infusion followed by isotopomer analysis via NMR or mass spectrometry [35]. In contrast, bolus administration of [U-^13^C_3_]pyruvate captures metabolic flux with minimal perturbation of endogenous metabolism. Across multiple pathways, ^13^C-labeled isotopologues showed temporal responses distinct from those of the corresponding unlabeled metabolite pools. In particular, alterations in labeled glutamate, citrate, succinate, and itaconate preceded or differed from changes in their endogenous pools, demonstrating earlier changes in metabolic flux than metabolite abundance. These findings extend previous hyperpolarized [1-^13^C]pyruvate studies reporting reduced [^13^C]bicarbonate and elevated [1-^13^C]lactate after TBI [11, 16, 20, 21, 39].

The metabolite-specific lesion patterns observed here also have potential translational implications. Composite metabolite ratios enhanced contrast between injured and intact tissue and integrated complementary aspects of neuroglial dysfunction and immunometabolism. Such spatial metabolic indices may therefore provide robust biomarkers for monitoring secondary injury and evaluating therapies targeting either neuroinflammation or metabolic dysfunction.

Rapid cryopreservation was selected to preserve both tissue morphology and metabolite localization for integrated MSI and IHC analyses. Although focused microwave fixation rapidly arrests enzymatic activity, intense and rapid tissue heating can complicate subsequent spatial metabolomic and immunohistochemical analyses, resulting in structural disruption and altered epitope preservation [10, 14]. For example, rapid heating of water-rich brain tissue can lead to vaporization-induced bubble formation, which may disrupt tissue architecture and compromise spatial integrity. Moreover, heterogeneous energy deposition may produce spatially uneven heating across the brain, particularly between cortical regions and deeper structures, potentially resulting in variable metabolic quenching. While rapid freezing cannot eliminate postmortem metabolic alterations, all specimens were processed using identical procedures, allowing reliable relative comparisons across brain regions and experimental time points.

## CONCLUSIONS

Our study demonstrates that TBI-induced immunometabolic remodeling is highly compartmentalized in both space and time. The integration of spatial metabolite mapping, immunohistochemistry, and *in vivo* ^13^C isotopomer tracing enabled characterization of neuroglial metabolic interactions during secondary brain injury and identifies candidate metabolic pathways and composite metabolic signatures that may facilitate the development of imaging biomarkers and ultimately guide therapeutic interventions targeting neuroinflammation and metabolic dysfunction.

## List of Abbreviations

CCI: controlled cortical impact
GFAP: glial fibrillary acidic protein
IHC: immunohistochemistry
MALDI: matrix-assisted laser desorption/ionization
MSI: mass spectrometry imaging
PC: pyruvate carboxylase
PDH: pyruvate dehydrogenase
RF: radiofrequency
ROI: region of interest
SDH: succinate dehydrogenase
TBI: traumatic brain injury
TCA: tricarboxylic acid
VIM: vimentin

## DECLARATIONS

## Ethics Approval and Consent to Participate

Not applicable.

## Consent for Publication

Not applicable.

## Availability of Data and Material

The MALDI MSI and IHC data were deposited to into the Mendeley Data and are available at the following URL: https://data.mendeley.com/datasets/mzbxchgszh/1

## Competing Interests

The authors declare that they have no competing interests.

## Funding

The National Institutes of Health of the United States (R01NS107409, P30DK127984, R01HL170039, S10OD028490), U.S. Army Medical Research Acquisition Activity (W81XWH2210485, HT94252510616), and Cancer Prevention & Research Institute of Texas (RP190617, RP240559, RP210099, RP260736).

## Authors Contributions

Conceptualization: JMP

Methodology: ZE, EHS, EJP

Investigation: ZE, EHS, BLBO, JMP

Data curation: ZE, EHS, JMP

Visualization: ZE, JMP

Funding acquisition: JMP

Project administration: ZE, JMP

Supervision: JMP

Formal analysis: ZE, JMP

Writing – original draft: ZE, EHS, EJP, JMP

Writing – review & editing: ZE, EHS, EJP, BLBO, JMP

## Acknowledgements

The CCI procedures were performed in the UT Southwestern Neuro-Models Facility (RRID:SCR_022529).

## Notes

### Competing Interest Statement

The authors have declared no competing interest.

