## Supporting Information for "Distinct spatiotemporal immunometabolic remodeling following acute traumatic brain injury"

### Supporting Figures

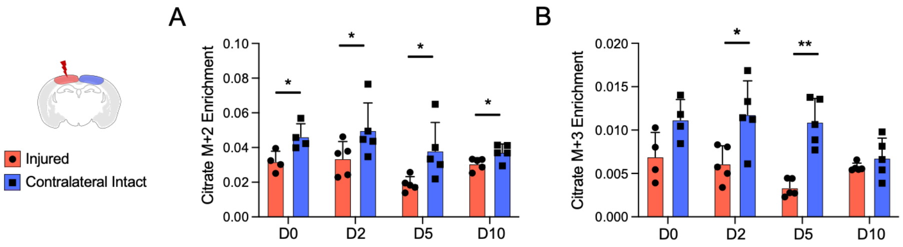


**Figure S1. Fractional enrichment of citrate isotopologues.**

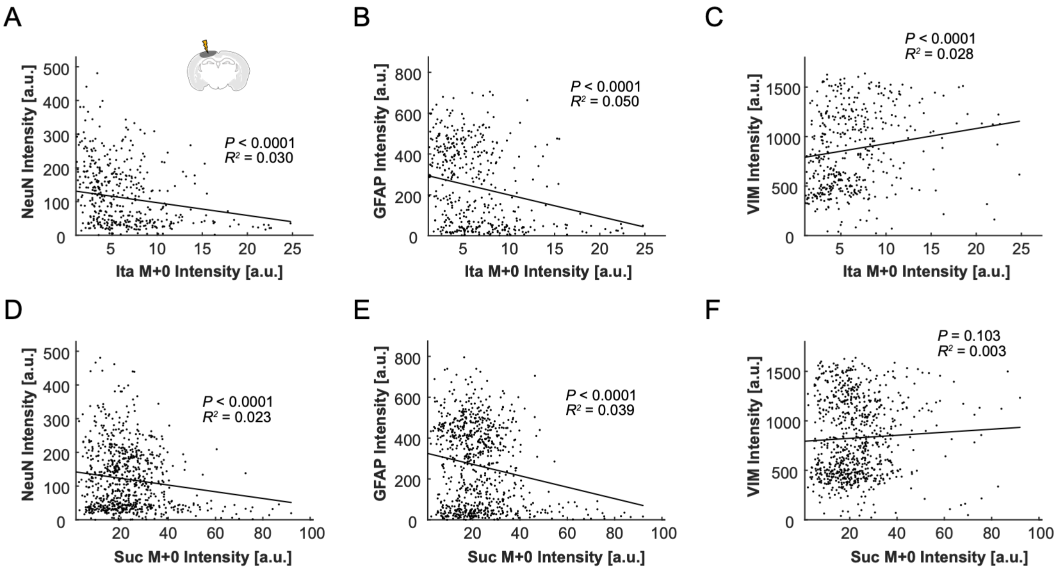


**Figure S2. Spatial correlation of itaconate and succinate with immunohistochemistry.** (A-C) Scatter plots of M+0 itaconate versus NeuN, GFAP, and VIM. (D-E) Scatter plots of M+0 succinate versus NeuN, GFAP, and VIM.

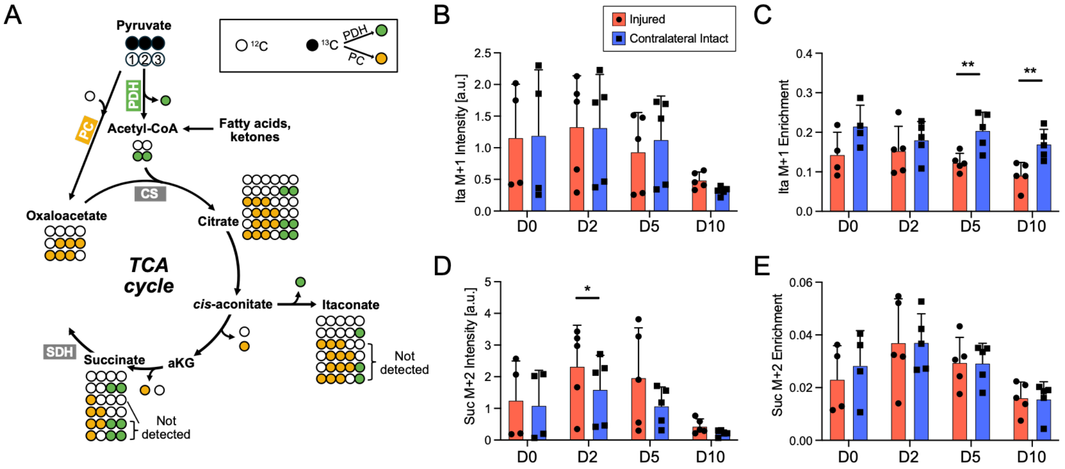


**Figure S3. Isotopologue analysis of itaconate and succinate.** (A) Schematic summarizing the metabolic origins of itaconate and succinate isotopologues. Pyruvate enters the TCA cycle via PDH (producing M+2 citrate) or PC (M+3 citrate), which give rise to downstream M+1 itaconate (via cis-aconitate) and M+2 succinate (via α-ketoglutarate). (B) M+1 itaconate and its (C) fractional enrichment. (D) M+2 succinate and (E) its fractional enrichment.

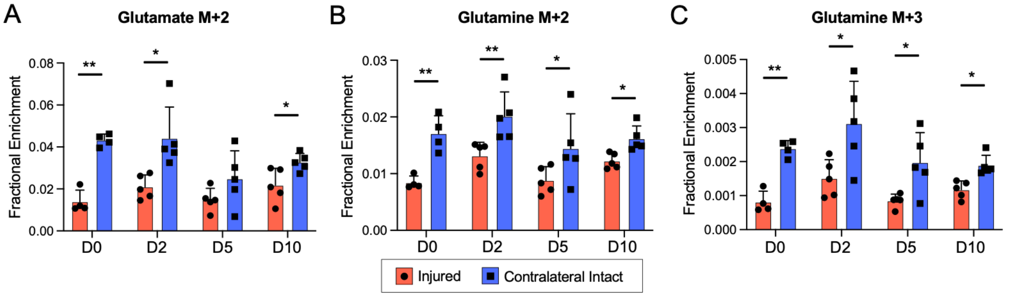


**Figure S4. Fractional enrichment of glutamate and glutamine isotopologues.**

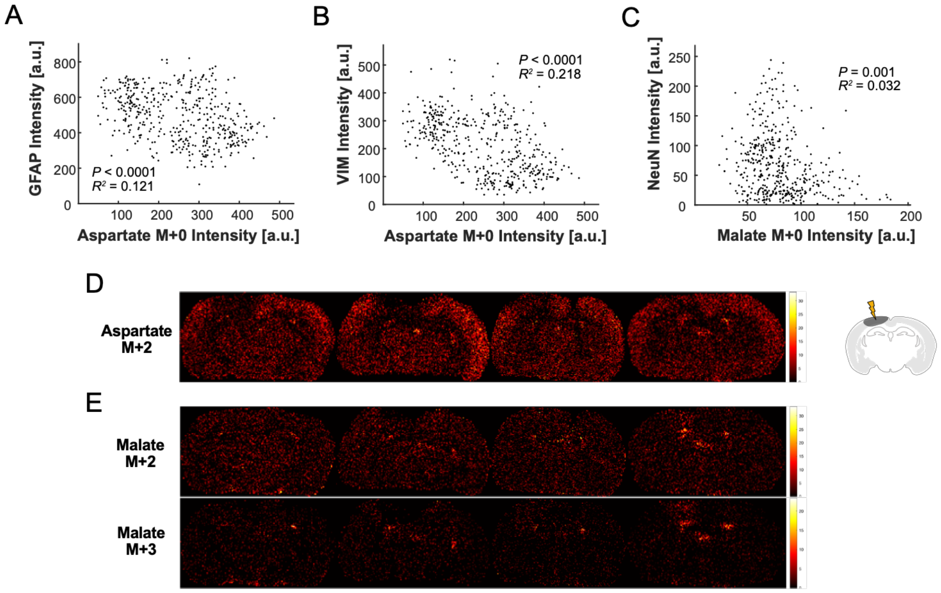


**Figure S5. Malate and aspartate isotopologue maps.** (A, B) Scatter plots of M+0 aspartate versus GFAP and VIM. (C) Scatter plot of M+0 malate versus VIM. (D) MALDI MSI of aspartate M+2. (E) MALDI MSI of malate M+2 and M+3.
